# Exploring trend and barriers of antenatal care utilization using data mining:evidence from EDHS of 2000 to 2016

**DOI:** 10.1101/351858

**Authors:** Kedir Hussein Abegaz

## Abstract

**Background:** The healthcare industry is paying attention to pregnancy and Antenatal care (ANC) for mothers. Thus, the presented study aimed at exploring the trend and identifying the barrier for ANC utilization of mothers in Ethiopia. Data mining is a field of big data science used to discover patterns and knowledge from big data.

**Methods:** All EDHS datasets from 2000 to 2016 were used for this study. The pooled cross-sectional study was conducted using the knowledge discovery process having steps; selection, cleaning, integration, transformation, and data mining algorithms. These algorithms are; Classification, clustering, association rules, and attribute ranking with pattern prediction.

**Results:** The proportion of ANC utilization was 27.6%, 28.2%, 34.5%, and 62.9% in 2000, 2005, 2011, and 2016 respectively. The pooled data; contained 28,631 mothers which were included in the study. Of these, more than half (56.09%) of them were not utilizing ANC during a pregnancy. Pregnancy complication, educational status of mothers and husbands, mothers’ residence, economic status, and media exposure had an association with ANC utilization having a confidence level of 95% and above.

**Conclusion:** ANC utilization in Ethiopia was increased significantly from 27.6%in 2000 to 62.9% in 2016. Despite this increment, the pooled proportion of ANC utilization is still low. The barriers to this low utilization were; Pregnancy complication, poor education of mothers and their husbands, living in rural, poor economic status, and media exposure. This study will recommend; firstly, pregnant mothers have to attend ANC service even though she had no pregnancy complication. Secondly, Education and poverty reduction are key strategic area to be addressed in improving women’s awareness towards ANC during a pregnancy. Thirdly, Expansion of infrastructure among the rural communities having good media coverage needs to be prioritized to improve ANC service utilization.

## Background

In the information industry, there are huge amounts of data available. These data are of no use till converted into useful information. Thus, making analyses and extracting useful information for this huge amount of data is essential. [1, 2]

The healthcare industry also contains massive amounts of data available [3]. Data generated in this sector includes relevant information about clients including their health care utilization status. The hidden trends, patterns, relationships and knowledge in healthcare data can be discovered using the application of data mining approaches [2, 4]. Data mining is a fast-growing field of big data science, sometimes known as knowledge discovery from a database (KDD).

Among the healthcare given to mothers, antenatal care (ANC) is the prominent care given to mother to identify, prevent, and treat conditions that may harm their lives and newborn babies [5, 6]. ANC is one of the four pillars of safe motherhood initiatives targeting the lessening of maternal and child mortality that can occur due to pregnancy-related complications [7].

Pregnancy-related deaths and diseases are remaining high globally in an unacceptable manner. In 2015, World Health Organization (WHO) estimated that 303,000 women died of pregnancy-related causes internationally. Developing countries shared 99% of these deaths and the countries in Sub-Saharan Africa (SSA) carried the largest burden of these maternal deaths responsible for 62%. This translates into an average maternal mortality ratio (MMR) of 510 deaths per 100,000 live births and a lifetime risk of 1 in 36 for a woman to die of maternal causes. Ethiopia is one of the SSA Countries with a high number of maternal mortality estimated that 13,000 in 2013. This is corresponding to an average MMR of 420 deaths per 100,000 live births. Among the top ten countries with high maternal mortality that shares 58% of the global maternal deaths; Ethiopia is the fourth in rank that is responsible for 4% of global maternal deaths annually [8].

The ample utilization of ANC service was found to be a significant contributor to the decrement of maternal mortality and morbidity [6]. Consequently, most countries including Ethiopia have developed and implemented national strategies towards safer motherhood by making ANC one of the major activities during a pregnancy [9]. According to the WHO recommendation [6] and current Ethiopian guideline on maternal care, ANC service has to be provided within four visits for women having normal progress on her pregnancy. The first visit is recommended being within the first trimester (16 weeks) of pregnancy, the second visit within 24-28 weeks, the third visit within 30-32 weeks and the fourth visit within 36-40 weeks of pregnancy [10]. Notwithstanding the availability of ANC services and the promising increment in the utilization in all health facilities of Ethiopia, the level of utilization is still low. During the five year period preceding Ethiopian demographic and health survey (EDHS) 2016, only 62% of women who gave birth during this period had at least one antenatal care (ANC) and only 32% of them had four or more ANC visits [11].

The previously done studies revealed that, ANC utilization was influenced by factors; Mother’s age [12–15], place of residence [16–19], women’s education [17, 20-22], husband education [19, 21], women empowerment [23], women’s occupation [24], economic status [13, 22, 25], parity[18, 24], history of pregnancy complication [18, 21], and media exposure [26, 27]. Even though these factors are from different studies, my study has included and attested all the variables.

The first purpose of this study is to look at the trend of ANC utilization in all EDHS. The second purpose is to identify the barriers to antenatal care utilization using data mining approaches based on the pooled EDHS. Hence, the results of this study have shown the evidence for programmers, decision makers, and respective stakeholders to use as a baseline in the efforts made to improve ANC services utilization in Ethiopia.

## Methods

### Study design and Data

The study design for the present study was pooled cross-sectional study and the data source was EDHS data of 2000, 2005, 2011, and 2016. These EDHS data were nationally representative at a particular time, and population-based cross-sectional surveys conducted in all regions of Ethiopia. The data were collected by the Central Statistical Agency (CSA) of Ethiopia in collaboration with International Classification of Functioning, Disability, and Health (ICF) and Measure DHS under the supports of the Ethiopian Ministry of Health. All mothers aged 15 to 49 and who had at least one live birth in five years preceding the surveys were eligible for this study.

The two-stage stratified cluster sampling with regions and residence as strata was used in all of the four surveys, EDHS 2000-2016. Enumeration Areas (EAs) from 1994 and 2007 population and housing census were the sampling units for the first stage sampling. In selecting EAs, 540 EAs for 2000 and 2005 surveys; 624 EAs to the 2011 survey; and 645 EAs to the 2016 survey were selected. Households (HHs) comprised the second stage of sampling. A representative sample of 11645 HHs in the 2000 and 2005 surveys, 17817 HHs in 2011, and 18008 HHs in the 2016 survey were selected. In my calculation, the total HHs number is 59,115. Of this HHs, 28,631 women who fulfilled eligibility criteria for the analysis of ANC services utilization were included. Women who had at least one live birth in five years preceding the surveys were eligible for this study.

### Knowledge discovery process (KDD)

KDD refers to the broad process of discovering knowledge in data and accentuates the “high-level” application of particular data mining methods. It is of interest to researchers in pattern recognition, machine learning, statistics, database, artificial intelligence, and data visualization [1].

Figure 1 presents the KDD for this study. In this figure, the KDD used to discover hidden and valuable knowledge from all EDHS (EDHS 2000-2016) datasets. The steps of this KDD include; Selection, Cleaning, Integration, Transformation, Mining.

**Figure 1:**
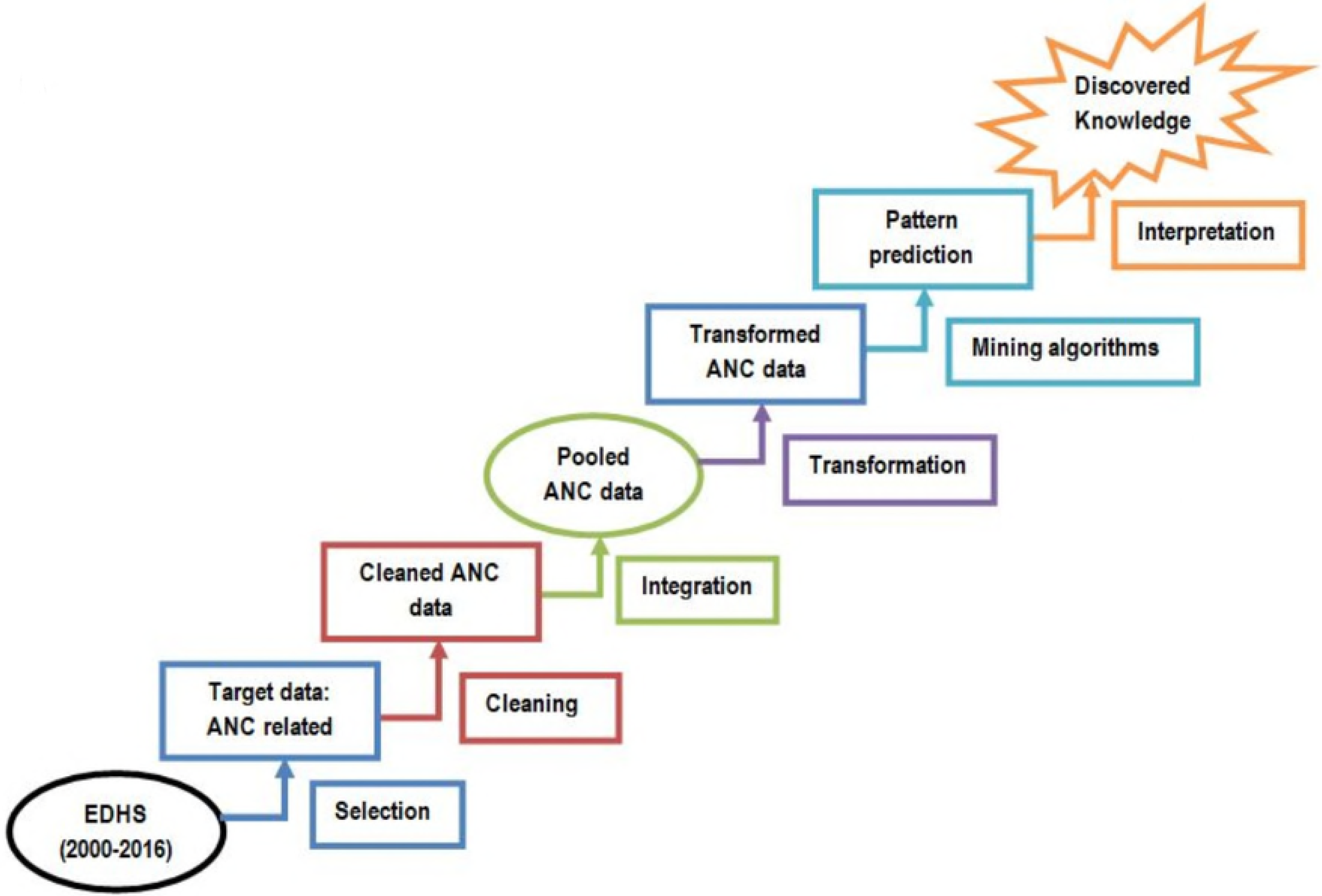
Knowledge discovery process of the presented study

### Data selection and cleaning

In the data selection step, a total of 10 attributes were selected, assuming to have a frequent pattern with ANC utilization based on the review from the previous literature dimensions in which ANC utilization was affected [12–27]. The dependent variable of this study is ANC utilization (“Yes” if the mother receives ANC for at least one time during a pregnancy, or else “No”).

The selected attributes were mother’s age, place of residence, mother’s education, husband’s education, woman’s empowerment (the empowerment to decide to go ANC), mother’s occupation, economic status, parity, history of pregnancy complications (if the mother had pregnancy complication at least once), and media exposure (the exposure to hearing media like TV and Radio,). Finally, the selected attribute is cleaned and recategorized to make suitable for data mining using STATA 14.

### Data integration and Transformation

All the EDHS data are integrated vertically depending on their similar attributes and a total of 28, 631 women were included in the analysis. In the transformation step, the integrated data was converted into “.arff” format which is appropriate to manipulate using a data mining tool.

### Data mining: Machine learning algorithms

The classification, clustering, and association and pattern prediction algorithms were applied to the pooled ANC data using the steps.

#### Classification algorithm

it extracts a model describing the important data classes [2]. It is one of the data mining techniques used to group the instances which belong to same class [2, 28]. Data classification has two steps; learning step (where a classification model is constructed) and classification step (where the model is used to predict class labels) for the given data [1]. The classification algorithms used in this study are; Tree structure (**ID3**), Statistical (**Naïve Bayes**), and Neural network (**Multilayer Perceptron**).

#### Clustering algorithm

clustering is the task of grouping a set of items in such a way those items in the same group (or cluster) are more similar to each other than to those in other groups. In clustering, it is required the users to input the number of clusters they desire as per the recommendation of most clustering algorithms [2, 29]. Thus, in this study, we have used five clusters using a center-based clustering **“Simple K-Means”** algorithm. It is based on the principle of maximizing intra-class similarity and minimizing inter-class similarity [1].

#### Association rules and Pattern prediction

Association rule learning is a rule based machine learning technique for discovering remarkable relations between attributes in the big database. It is anticipated identifying strong rules discovered in databases using some measures of interestingness [30].

Finding the frequent itemsets from a given dataset (ranking) and deriving association rules are among the most popular data mining approaches [31]. Finding frequent item-sets is important because of its combinatorial explosion. If frequent item-sets are found once, it is straightforward to generate association rules with confidence larger than or equal to a user-specified minimum confidence.

For this study, the **“Apriori algorithm”** (a seminal algorithm for finding frequent itemsets using candidate generation) [31, 32] is used to identify the best associations rules and **“InfoGainAttributeEval”** with **“Ranker-T”** to rank the attributes of the pooled ANC dataset. All the analyses were explored using the well-known and open source data mining tool WEKA 3.7 [33].

## Results

A total of 28, 631 mothers were included in this study. **Table 1** presented the summary statistics of ANC utilization of these mothers from 2000-2016. This result revealed that more than half of mothers (56.1%) were not utilizing ANC, a great majority of mothers (81.5%) were from the rural area of residence, majority (70.3%) of them and their husband (53.4%) were illiterate, nearly half (46.9%) of them were poor economically. Finally, More than 65% of mothers were empowered fully and partially.

**Table 1:** Summery statistics and association between ANC Utilization and independent variables

| Variables with category |  | Freq. (%) | Prob.<br>Pearson chi<sup>2</sup> |
| --- | --- | --- | --- |
| ANC Utilization | ANC Utilized | 12,571 (43.91) |  |
|  | ANC not utilized | 16,060 (56.1) |  |
| Women empowerment | Full empowered | 3,269 (15.2 ) | 0.000 |
|  | Partially empowered | 10,970 (51.1 ) |  |
|  | Not empowered | 7,214 (33.6) |  |
| Media Exposure | No exposure | 18,830 (65.8) | 0.000 |
|  | Rare Exposure | 8,526 (29.8) |  |
|  | Frequent exposure | 1,263 (4.4) |  |
| Place of residence | Urban | 5,293 (18.5) | 0.000 |
|  | Rural | 23,338 (81.5) |  |
| Mother's Education | No education | 20,132 (70.3) | 0.000 |
|  | 1<sup>o</sup> education | 5,998 (21.0) |  |
|  | 2<sup>o</sup> and higher | 2,501 (8.7) |  |
| Economic status | Poor | 10,056 (46.9) | 0.000 |
|  | Medium | 3,438 (16.0) |  |
|  | Rich | 7,959 (37.1) |  |
| Number of children ever born | 1 | 5,525 (19.3) | 0.000 |
|  | 2-4 | 12,248 (42.8) |  |
|  | 5 and more | 10,858 (37.9) |  |
| Age of women | Less than 25 | 1,636 (5.7) | 0.000 |
|  | 25-34 | 19,644 (68.6) |  |
|  | 35-49 | 7,351 (25.7) |  |
| Pregnancy complication history | Had history | 4,552 (36.2) | - |
|  | Had no history | 8,019 (63.8) |  |
| Husband's education | No education | 15,278 (53.4) | 0.000 |
|  | 1<sup>o</sup> education | 8,065 (28.2) |  |
|  | 2<sup>o</sup> and higher | 5,288 (18.5) |  |
| Mother's Occupation | Had no job | 15,160 (53.0) | 0.000 |
|  | Had job | 13,471 (47.1) |  |

The association between ANC utilization and other variables are also presented in Table1. This result implies that all the variables in this table had an association, supported with the Pearson chi-square p-value of less than 0.05. Therefore, these variables are candidate variables to be tested as barrier factors to the mother for their ANC utilization.

The three classification algorithms for classification of instances in the pooled data were Decision tree (Id3), Naïve Bayes, and Multilayer Perceptron. Based on this classification result, Decision tree classified instances correctly with the highest precision, 73.3%. This is to mean that, from mothers classified as ANC Utilized, only 73.3% of the were correctly classified as ANC utilized but 26.7% were incorrectly classified as ANC utilized.

In the trend of ANC utilization of mothers, the proportion of ANC utilization was 27.6% in EDHS 2000, 28.2% in EDHS 2005, 34.5% in EDHS 2011, and 62.9% in EDHS 2016 respectively. This result showed that the ANC utilization was increased from EDHS 2000 to EDHS 2016. This increment proofs that the ANC Utilization in Ethiopia was improved from time to time. The possible reasons for this increment will be, the expanding of health care services to rural areas of the country, the awareness of mothers towards ANC was in betterment from year to year.

The clustering of mothers regarding their ANC utilization was displayed in Table 2. In this result, mothers were clustered into five groups; 1^st^, 2^nd^ 3^rd^, 4^th^, and 5^th^ cluster. One third of mothers (33.5%) were grouped in a 2^nd^ cluster with the following characteristics; ANC not utilized, had no history of pregnancy complication, no mother education, rural place of residence, no husband education, economically poor, partially empowered, having no media exposure, had no job, having 2-4 children, and their age was from 25-34. In reverse of this, the 4^th^ cluster showed that mothers who were utilizing ANC, had history of pregnancy complication, had secondary and higher educational status of mothers and husbands respectively, were living in urban, rich economically, partially empowered, had rare exposure of media, had a job, had only 1 child, and their age was from 25-34. This result revealed that mothers having the above-listed characters were utilizing ANC.

**Table 2:** Clusters of mothers based on their similar characteristics

| Attribute | 1<sup>st</sup> Cluster<br>(32.5%) | 2<sup>nd</sup> Cluster<br>(33.5%) | 3<sup>rd</sup> Cluster<br>(11%) | 4<sup>th</sup> Cluster<br>(14%) | 5<sup>th</sup> Cluster<br>(8%) |
| --- | --- | --- | --- | --- | --- |
| ANC utilization | Not utilized | Not utilized | No utilized | Utilized | Not utilized |
| Mother education | No education | No education | Primary | 2<sup>o</sup> & higher | Primary |
| Place of Residence | Rural | Rural | Rural | Urban | Rural |
| Husband education | No education | No education | Primary | 2<sup>o</sup> & higher | Primary |
| Economic status | Poor | Poor | Rich | Rich | Poor |
| Women Empowerment | Partial empowerment | No empowerment | Partial empowerment | Partial empowerment | Partial empowerment |
| Media exposure | No exposure | No exposure | No exposure | Rare exposure | No exposure |
| Mother occupation | Had no job | Had no job | Had no job | Had job | Had job |
| Parity | 2-4 | > =5 | > =5 | 1 | 1 |
| Mother's age | 25-34 | 25-34 | 35-49 | 25-34 | 25-34 |

Table 3 displayed the top five association rules in predicting the important attributes of ANC utilization. In this result, the maximum confidence attained was 100% from the first rule. This rule implies that all mothers who had pregnancy complication were utilized ANC service. The second, third, fourth and fifth rules had a confidence level of 95% and above. The second rule implies if the mother had no education and she was poor economically, she had no ANC utilization. The third rule displays that, if the mother had no media exposure and she was poor economically, she was not utilizing ANC. The fourth rule tells us, if the husband had no education and she was living in a rural area, she was not utilizing ANC. Finally, the fifth rule displays that, if the mother was living in rural area and she had no media exposure, she was not utilizing ANC.

**Table 3:** The top five association rules in predicting barriers for ANC Utilization

| Rank | Association Rules | Conf. level |
| --- | --- | --- |
| 1<sup>st</sup> | {Preg_comp=No}→{ANC utilization=No} | conf:(1) |
| 2<sup>nd</sup> | {Mother_Edu=No Education & Econ_stat=Poor }→{ANC utilization=No} | conf:(0.99) |
| 3<sup>rd</sup> | {Media_exp=No exposure & Econ_stat=Poor}→{ANC utilization=No} | conf:(0.98) |
| 4<sup>th</sup> | {Husb_Edu=No Education & Residence=Rural} →{ANC utilization=No} | conf:(0.96) |
| 5<sup>th</sup> | {Residence=Rural & Media_exp= No exposure}→{ANC utilization=No} | conf:(0.95) |

Table 3 displayed also, pregnancy complication, educational status of mothers, economic status, media exposure, husband education, and mothers’ residence were important attributes for ANC utilization having a confidence level of 95% and above. This means that the above-listed variables in this table had an association to be important barriers for ANC utilization.

Figure 2 presented patterns for predicting the barriers of ANC utilization; this was done by identifying the most frequently occurred attributes. This result shows that among the 10 attributes included in this study; Pregnancy complication was the first most frequently occurred attribute in all patterns, Mothers’ education, Place of residence, Husband education, Economic status and Media exposure were second, third, fourth, fifth, and sixth frequent attributes to occur.

**Figure 2:**
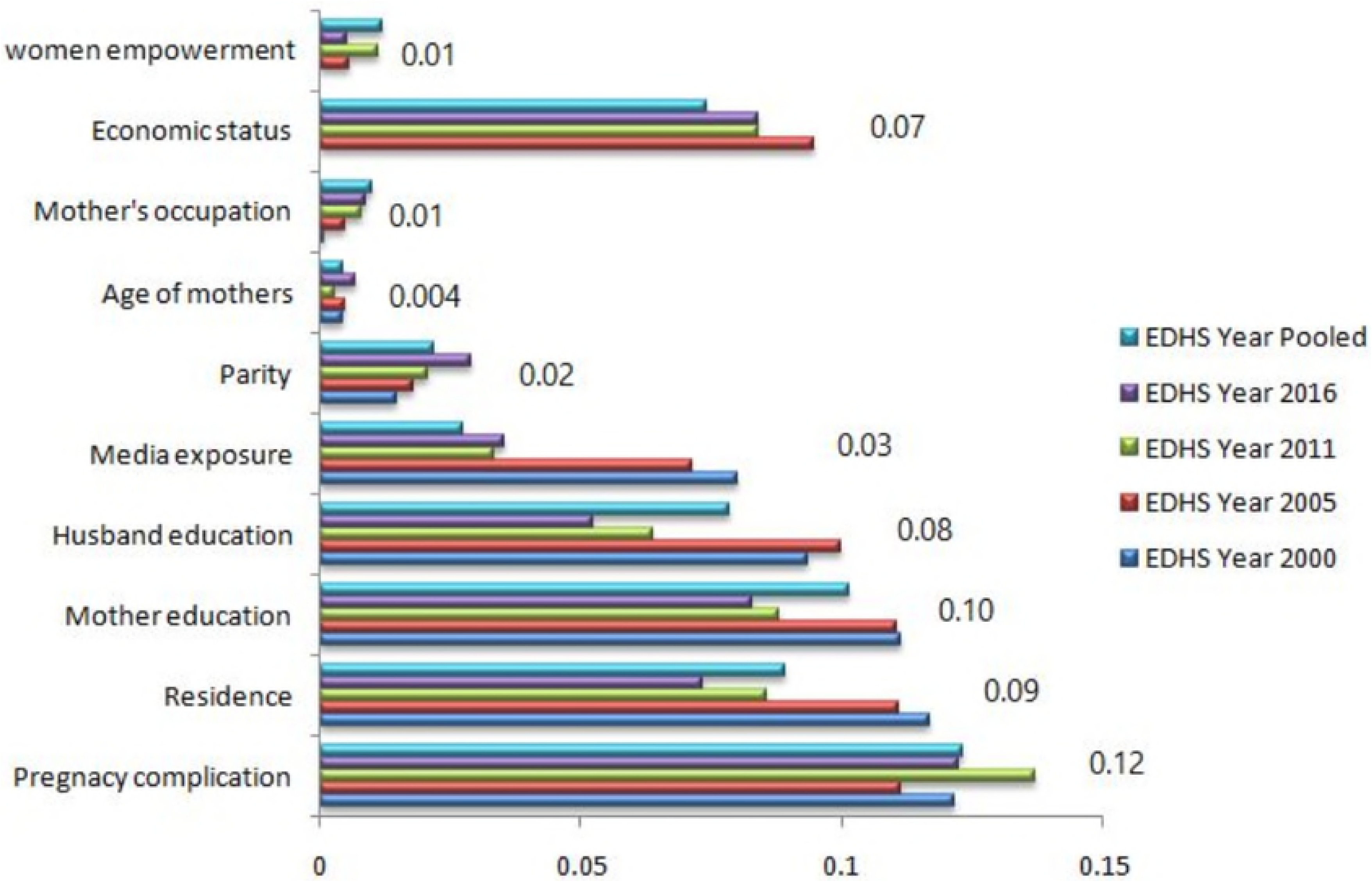
Identified patterns for barriers of ANC Utilization

These results in pattern prediction and association rule suggest that, among all 10 attributes having an association, pregnancy complication, mother’s and husband’s educational status, place of residence, economic status, and media exposure were the predictors for barriers of ANC utilization.

## Discussion

The presented study aimed to determine the trend and barriers of ANC utilization of mothers in Ethiopia. The current study revealed that the trend of ANC utilization was increased from EDHS 2000 to EDHS 2016, with a pooled proportion of only 43.9% of mothers were utilized ANC. Ethiopia is among countries achieved the lowest proportion of ANC utilization compared to countries like Zambia and Cameron [34, 35]. This low proportion of ANC utilization might be due to the reason that parallel to the strategic interventions to improve ANC service utilization coverage, the adequacy of care has not been prioritized enough and addressed as the strategic area in Ethiopia. In this study, the likelihood of ANC utilization was associated with barriers like pregnancy complications; mothers and husbands lower educational status, poor economic status, place of residence, and had no media exposure.

The presented study elucidated that, the utilization of ANC service was strongly influenced by mother’s history of pregnancy complications. When mothers had a history of pregnancy complications, they were attending ANC service. This is agreed with the previous findings of studies that undertaken in Ecuador and Taiwan [18, 21]. This is probably due to those women who experienced pregnancy complications before are more concerned about their health and better perceived the risk of pregnancy. As a result, they are more probably keen to seek medical care early and regularly.

The current study showed that the chance of ANC utilization was positively affected by women’s level of education. Similar discoveries were exposed in previous studies that undertaken in Vietnam, Taiwan, and India [17, 21, 22]. This finding might be due to women education does have the tendency too proximally affect their improvement in other socioeconomic and socio-demographic aspects. Women education could also support their financial autonomy in traveling to have a medical care, their exposure status to media as well as access to information and their ability to understand easily and positively evaluate the trade-off either side of the care. In the present study, husband education was also positively affected women to utilize ANC service. Previously conducted studies in Nepalese and Taiwan support this finding [19, 21]. This is because the educated husbands are more likely to have awareness of the importance of health care and more likely to support women in having care contrary to those with no education.

Household wealth index also made an impact on the utilization of ANC. As per this finding, the likelihood of using ANC increased along with the change in the household wealth index from poor to rich. This finding was revealed in previous studies that were conducted in Brazil, India and other developing countries [13, 22, 25]. In fact, Ethiopia has declared ANC services to be provided for free to all mothers in order to avoid the financial barrier those women could encounter while seeking the care. However, this alone could not totally avoid the financial barriers to seeking health care during a pregnancy. The cost that comes along with traveling to reach distant health institutions could hinder the need and make women reluctant for an early initiation and subsequent visits. Mostly they don’t travel alone to a distant health facility, rather either with their husbands or children. Affording for traveling and other related costs may be another barrier even if the services are provided for free.

Frequent media exposure like; Television, Radio, and Magazine had a positive influence on women to utilize ANC service. Previously conducted studies in Ethiopia and Nigeria witnessed the influence of media exposure on ANC visits [26, 27]. The frequent promotion of maternal health services through media could influence women’s predisposition for an early visit and their adherence to subsequent follow-ups by providing them with relevant information concerning the risk of pregnancy and the benefit of services.

Being in a rural area was among prominent barriers to ANC utilization. The chance of using ANC services was considerably reduced among women from the urban community to women from the rural community. This result was consistent with the finding of studies done in Vietnam, Ecuador and Nepalese [17–19]. This is most probably due to many social infrastructures; including health, education, transport, and information are highly concentrated in urban areas compared to rural areas. The best availability of these infrastructures in urban areas may have an important role in supporting women to develop a good health care seeking behaviors.

## Conclusion

The proportion of ANC utilization of EDHS was increased significantly from 2000 to 2016. Had pregnancy complication, low education of mothers, Low economic status, no media exposure, poor husband education, the rural residence of mothers were the barriers predicted for the utilization of ANC.

### Recommendations

Despite the availability of maternal health care in Ethiopia, the level of ANC utilization is still low. In the presented study, it was observed that pregnancy complication was one of the prominent attributes. Then, mothers have to be aware that, ANC utilization service is not only for mothers having a complication. Therefore, the first recommendation will be, pregnant mothers have to attend ANC service even though she had no pregnancy complication. Mother’s and husband’s education was the important attributes for ANC service utilization in Ethiopia. Therefore the second recommendation will be education is a key strategic area to be addressed by the ministry of education of Ethiopian in improving women’s awareness towards ANC during a pregnancy. Along with this, the economic disparity is the influential attributes that contributed to the difference among women in using ANC services. Therefore, the third recommendation will be poverty reduction, which is another area of intervention that needs to be addressed by the Ethiopian government. The urban women were substantially different from using ANC services from the rural community. Along with this, perceived distance problem is a barrier that impedes women from utilizing adequate ANC. Therefore, the fourth recommendation will be the expansion of infrastructure among the rural community needs to be prioritized by the Ethiopian government, to improve ANC service utilization. Media exposure is also among the important attributes for ANC utilization. Finally, this study will recommend that the government of Ethiopia has to expand the media coverage related to ANC throughout the country including, and mothers have to be aware of the importance of ANC during pregnancy.

## List of Abbreviations

ANC: Antenatal Care
CSA: Central Statistical Agency
DHS: Demographic and Health Survey
EAs: Enumeration Areas
EDHS: Ethiopian Demographic and Health Survey
ICF: International Classification of Functioning, Disability, and Health
KDD: Knowledge Discovery from Database
MLP: Multilayer Perceptron
MMR: Maternal Mortality Ratio
SSA: Sub Saharan Africa
WEKA: Waikato Environment for Knowledge Analysis
WHO: World Health Organization

## Declarations

### Ethical Approval and Consent to participate

Ethics approval and participant consent were not necessary as this study involved the use of a previously-published de-identified database by Central Statistical Agency of Ethiopia.

### Consent for publication

Not Applicable

### Availability of supporting data

The dataset was demanded and retrieved from DHS website https://dhsprogram.com after formal online registration and submission of the project title and detail project description.

### Authors’ Contribution

KHA performed the analysis, wrote and approved the final manuscript.

## Acknowledgments

None

## Authors’ Information

Biostatistics and Health Informatics, Public Health Department, Madda Walabu University, Ethiopia

